# Antigenic and genetic diversity of H1 and H3 influenza A viruses in swine in Brazil

**DOI:** 10.1101/2023.12.01.569635

**Authors:** Sara Lopes, Tavis K. Anderson, Rejane Schaefer, Caroline Tochetto, Danielle Gava, Mauricio E. Cantao, Janice R. Ciacci-Zanella, Amy L. Vincent Baker, Nicola S. Lewis

**Affiliations:** Department of Pathobiology and Population Sciences, The Royal Veterinary College, University of London, Hertfordshire, UK; Virus and Prion Research Unit, National Animal Disease Center, USDA-ARS, Ames, IA, USA; Embrapa Suínos e Aves, BR 153, Km 110, Distrito de Tamanduá, Concordia 89715-899, SC, Brazil; Worldwide Influenza Centre, The Francis Crick Institute, London, United Kingdom

**Author notes:** Address correspondence to: Rejane Schaefer, Embrapa Suínos e Aves, BR 153, Km 110, Distrito de Tamanduá, Concordia 89715-899, SC, Brazil., Nicola S. Lewis, Department of Pathobiology and Population Sciences, The Royal Veterinary College, Hawkshead Lane, North Mymms, Hatfield, Hertfordshire, AL9 7TA, United Kingdom.

**Keywords:** Influenza A, Surveillance, Antigenic diversity, Swine, Vaccine

## Abstract

**Background:** Influenza A virus (IAV) circulates within human and swine populations, and pigs are considered intermediate hosts for the generation of IAV with pandemic potential. Surveillance and characterization of IAVs circulating in pig populations are crucial to strain match vaccines to control IAV transmission in pigs and quantify pandemic potential to humans.

**Methods:** Here, we characterized the genetic and antigenic diversity of IAVs circulating in Brazilian swine between 2010-2018. Phylogenetic maximum-likelihood trees were generated for 84 Brazilian hemagglutinin (HA) gene segments. Hemagglutination inhibition (HI) assay data was used with antigenic cartography to quantify the antigenic differences among representative H1 and H3 swine viruses and relative cross-reactivity between these viruses and human seasonal vaccine strains.

**Results:** We identified two genetic lineages of H1 viruses derived from separate human-to-swine transmission events (H1 1B lineage, clades 1B.2.3 and 1B.2.4), an H3 lineage that has diversified into two genetic clades (H3 1990.5.1 and 1990.5.2), and HA genes associated with the 2009 H1N1 pandemic. There was limited cross-reactivity between circulating swine lineages and significant antigenic variation within lineage.

**Conclusions:** The antigenic diversity among endemic IAV in swine indicates a need for regional strain-specific vaccination strategies in Brazil. Our data supports the need for systematic genomic surveillance and characterization in Brazil to improve the efficacy of swine vaccines and quantify the pandemic potential of endemic swine influenza A viruses.

## 1. Introduction

H1 and H3 subtype influenza A viruses (IAV) circulate in both human and swine populations, with occasional human-to-swine transmission events increasing the genetic and antigenic diversity of IAV within pigs (1, 2). Swine-to-human events also occur, causing sporadic limited zoonotic infections and rare outbreaks or pandemics. A novel IAV of swine origin, A(H1N1)pdm09, emerged in Mexico in early 2009 and, spread rapidly worldwide, causing the first pandemic of the 21^st^ century (3–5). In Brazil the first report of A(H1N1)pdm09 in swine was in January of 2010 and several outbreaks were subsequently reported in regions of Brazil where there is extensive pig production (6).

Brazil is the fourth largest pork producer and exporter in the world; however, there are few reports describing circulation of IAVs in Brazilian swine populations prior to the 2009 pandemic (6, 7). Previous work identified multiple human-to-swine spill-overs of seasonal H3 and H1 influenza A virus strains into pigs in Brazil (8, 9). Genes from these spillovers persisted in Brazilian swine and the internal gene cassette of these viruses has changed through reassortment with the A(H1N1)pdm09 that now also co-circulates within the swine population (9, 10). The impact of reassortment and cocirculating lineages of viruses on antigenic diversity in Brazilian pigs is unresolved. Vaccination can be used to mitigate influenza A disease risk in pigs. However, for maximum efficacy, vaccine strains should match with currently circulating viruses and this requires periodic assessment of the antigenic and genetic relationships between circulating strains and vaccine strains (11, 12). Vaccine strains with significant antigenic drift from circulating viruses may not confer sufficient protection against future infections on farms. In Brazil there is only one commercial H1N1pdm vaccine available (FluSure® by Zoetis) and only one laboratory in the country produces autogenous vaccines for IAV.

In this study, we characterized the antigenic diversity of IAV strains in Brazilian swine isolated between 2010-2018 and the relationship to human seasonal vaccine strains. We integrated antigenic and genetic data and quantified the relationship among strains using antigenic cartography to determine the potential molecular basis for the antigenic variation and virus evolution of IAV within Brazilian swine. These data form an objective basis to select vaccine components better matched to circulating swine IAVs in Brazil and assess the potential zoonotic risk of circulating strains in Brazilian pigs.

## 2. Materials and Methods

### 2.1 Virus sampling, diagnostic testing and sequencing

Nasal swabs and lung samples were collected from commercial swine in herds located in six Brazilian states and sent to a diagnostic laboratory for screening for respiratory pathogens involved in the porcine respiratory disease complex. Samples positive for IAV by RT-PCR (13) were submitted for virus isolation in SPF chicken eggs and/or MDCK cells (14) and virus isolates were sequenced. Briefly, RNA was extracted from virus isolates and the eight influenza gene segments were amplified by RT-PCR using PathAmp FluA reagents. DNA libraries were prepared and submitted for sequencing using Ion Torrent system. Influenza genomes were assembled using Newbler V 2.9. Geographical maps with the location where each Brazilian virus was detected and isolated were generated using MapChart.

### 2.2 Phylogenetic and Genetic Analysis

A total of eighty-four (84) Brazilian hemagglutinin (HA) gene sequences were generated and an additional 499 randomly sampled sequences from the same time period (2010 to 2018) were retrieved from the Influenza Research Database (15). The final dataset of 508 H1 sequences and 269 H3 sequences also contained reference World Health Organization (WHO)-recommended human seasonal H3 and H1 HA vaccine sequences and candidate vaccine sequences (sequence accessions and input alignments are available at https://github.com/flu-crew/datasets).

Sequence alignments were constructed for H1 and H3 HA gene segments with default settings in MAFFT version 7.222 (16). The segments were trimmed to only retain nucleotides from the starting ATG until the final stop codon. We inferred Maximum Likelihood (ML) phylogenetic trees for each of the segments using IQ-TREE, v.1.5.5 (17) using the automatic model selection process (18) and statistical support determined using the ultrafast bootstrap approach and the SH-like approximate likelihood ratio test (19, 20). From the inferred phylogenetic trees, clades were defined based on the sequences sharing a common node, monophyly, and statistical support (ultrafast bootstrap >70, aLRT support >90). To produce a reference panel for antigenic characterization we selected one virus from each circulating clade in Brazil in order to raise swine antisera. To span genetic diversity for each clade, an HA1 amino acid consensus sequence was determined, and matching field strains were selected as a representative for each major clade of the H1 and H3 within the phylogenetic tree and used as a test antigen for subsequent hemagglutination inhibition (HI) assays. A similar analytical process was undertaken to select representative viruses to raise reference antisera for HI assays. Clade classifications for the Brazilian H1 and H3 HA genes were assessed using the Swine H1 Clade Classification tool (21) and the octoFLU classifier (22).

### 2.3 Amino Acid Comparison Analysis

Amino acid differences for the HA1 genes within each of the defined clades were calculated using custom python script (https://github.com/flu-crew/flutile). We compared the isolated Brazilian viruses with their closest ancestral and more current human influenza vaccine strains. The amino acid sequences of Brazilian H1 1A viruses were compared with 1A.3.3.2 A/Michigan/45/2015, an A(H1N1) pdm09 virus included in the composition of the 2018-2019 seasonal influenza vaccine. Brazilian H1 1B.2.3 and 1B.2.4 clade genetic sequences were compared to human seasonal vaccine H1 clade strains A/Solomon Islands/3/2006 and A/New Caledonia/20/1999, which were the closest seasonal influenza vaccine recommendation to each ancestral introduction into Brazilian pigs. Amino acid positions in the Brazilian H3 sequences were numbered according to the Burke and Smith numbering system (23) and H3 sequences were compared to human seasonal vaccine H3 strain A/Wuhan/359/1995. HA gene segments from Brazilian strains were compared for amino acid differences. Differences located in putative antigenic sites (24, 25) that have been previously identified to cause antigenic change in influenza A viruses were annotated.

### 2.4 Hemagglutination Inhibition Assay (HI) and Antigenic Cartography

Monovalent swine antisera were generated from representative swine IAV and were heat inactivated at 56°C for 30min. Hemagglutination inhibition assays (HI) were conducted against a panel of global strains of swine and human IAVs using turkey red blood cells (26). We merged this HI dataset with previously generated and unpublished H1 and H3 HI assay data from United States Department of Agriculture (27–29).

HI data were used to generate 3-dimensional (3D) antigenic maps to quantify the antigenic relationships among the Brazilian swine IAVs and the global panel using antigenic cartography (30). Antigens that showed less than 3 numeric titers across the HI assay were removed as these were not robustly positioned in the maps. Antigenic maps were generated in RStudio 1.1.442 (31) and Acmacs Online Software (https://acmacs-web.antigenic-cartography.org/). The antigenic distances between the Brazilian H1 and H3 strains in pigs and the reference human seasonal strains were extracted and plotted in RStudio 1.1.442. The distribution of the distance data was illustrated in a boxplot with the median of the data shown as a line and the bottom and the top of the rectangle representing the lower quartile (Q1/25% of the data) and upper quartile (Q3/75% of the data) respectively. The whiskers attached to the rectangle indicate variability with the Q1 and Q3 quartiles and points positioned outside of the whiskers of the boxplot are outlier values.

### 2.5 Identifying the molecular determinants of antigenic diversity

To explore the molecular basis to the observed antigenic diversity in Brazilian pigs we first compared the inferred amino acid sequences of the HA1 protein of the Brazilian viruses and identified amino acid substitutions in previously identified antigenic sites, which also defined the antigenic groups seen in the antigenic maps. We compared the HA1 genes of the 1A.3.3.2 Brazilian viruses against A/Michigan/45/2015, the vaccine strain that represented the closest ancestor A(H1N1)pdm09 strain to identify the amino acid differences between the putative ancestor and current strains that might account for the antigenic variation since introduction from humans. A similar analysis was also conducted for the 1B strains where we generated amino acid sequences for the HA1, and compared the swine viruses against the most similar human seasonal vaccine in our dataset (human seasonal vaccine A/Solomon Islands/3/2006 strain, or A/New Caledonia/20/1999). For the H3 swine viruses, we generated HA1 amino acid sequences and compared these against the human seasonal vaccine A/Wuhan/359/1995, and identified diversity within known epitope positions (24) and the previously described “antigenic motif” defined using H3 viruses circulating in US swine (27, 32).

## 3 Results

### 3.1 Genetic evolution of swine influenza A viruses in Brazilian pigs

In the global phylogenetic trees, the sequence data described here and used in antigenic analyses included 36 H1N1 viruses from Brazilian pigs that were genetically related to A(H1N1)pdm09 strains from the H1 1A.3.3.2 lineage (Figure 1). There were at least 8 human-to-swine transmission events that occurred from 2010 to 2018 and were previously described (8). Of these humans to pig introductions, one resulted in sustained transmission of the virus in Brazilian swine populations with 24 subsequent detections in this time period that have been genetically characterized in detail (10).

**Figure 1.**
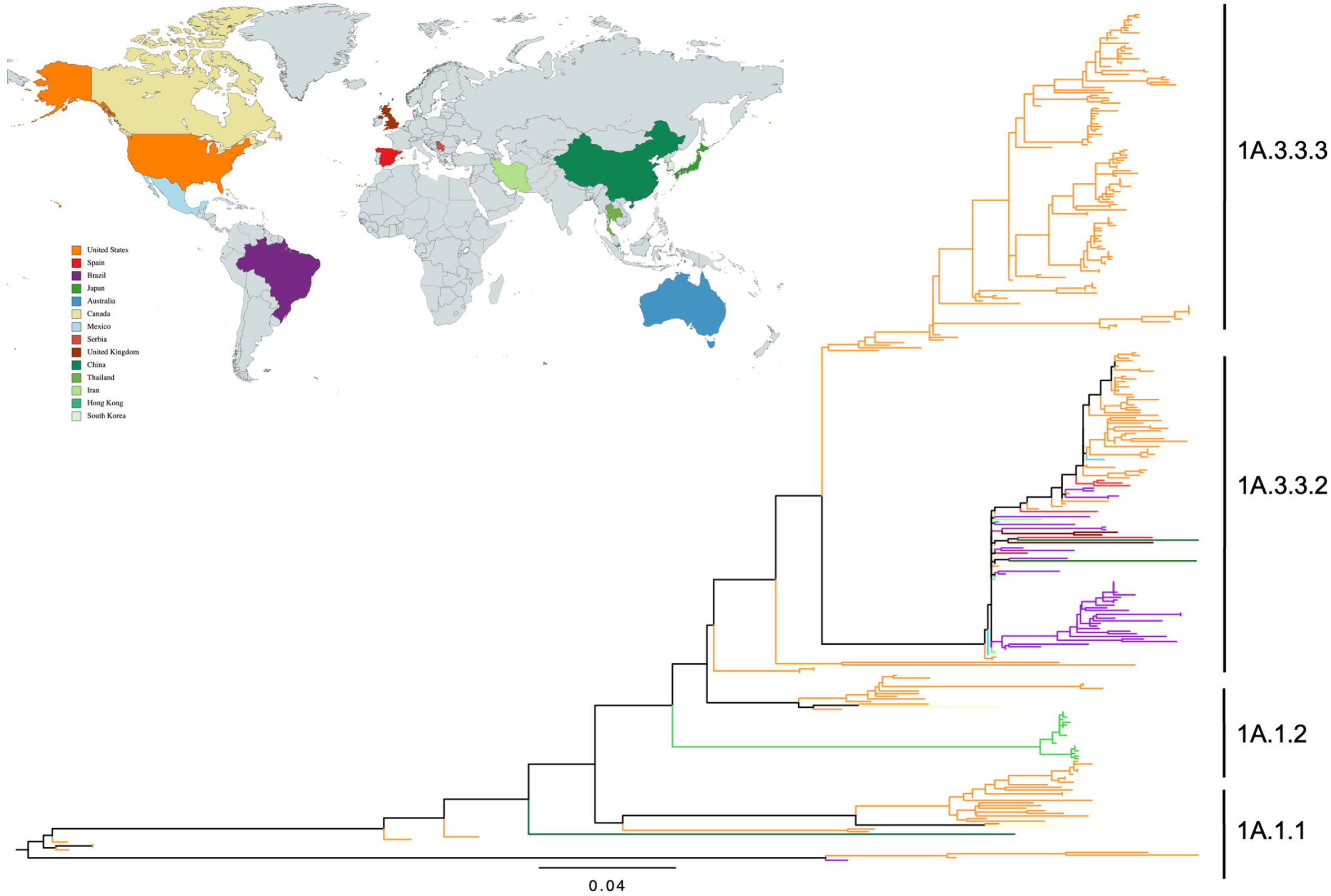
Phylogeny and geographic distribution of H1 1A classical swine lineage influenza A virus in swine hemagglutinin genes collected in Brazil. Mid-point rooted maximum-likelihood tree inferred for Brazilian swine HA gene sequences detected and isolated between 2010 and 2018 and global swine isolates, human seasonal strains, human vaccine, and human variant HA genes. The tree branches were colored by geographic isolation of the different strains and Brazilian HA genes were colored purple and the geographical map was generated using MapChart.

Twenty-three H1N2 and 2 H1N1 strains from Brazilian pigs were genetically related to human-like H1 1B lineage strains (Figure 2). The H1N2 strains currently circulating in Brazil were the result of at least two separate introductions from humans in the early 2000s supporting prior research (8). The genetic diversity, statistical support, and evidence of onward transmission of these clades observed in this study warrant the development of two new clade designations within the 1B lineage; 1B.2.3 and 1B.2.4 (following Anderson et al. (21)). The 1B.2.4 phylogenetic clade was detected more frequently and has diversified since the introduction in the early 2000s to consist of three statistically supported clades. The human-to-swine introduction that established the 1B.2.3 clade was first detected in 2011 and also evolved post-introduction. Both 1A and 1B lineages co-circulated in Brazilian pig farms at least since 2011. 1A lineage strains were also detected in Rio Grande do Sul, Santa Catarina, Minas Gerais and Parana (Figure 4). 1B lineage clades were detected in Santa Catarina, Rio Grande do Sul, Parana, Minas Gerais and Mato Grosso do Sul (Figure 5).

**Figure 2.**
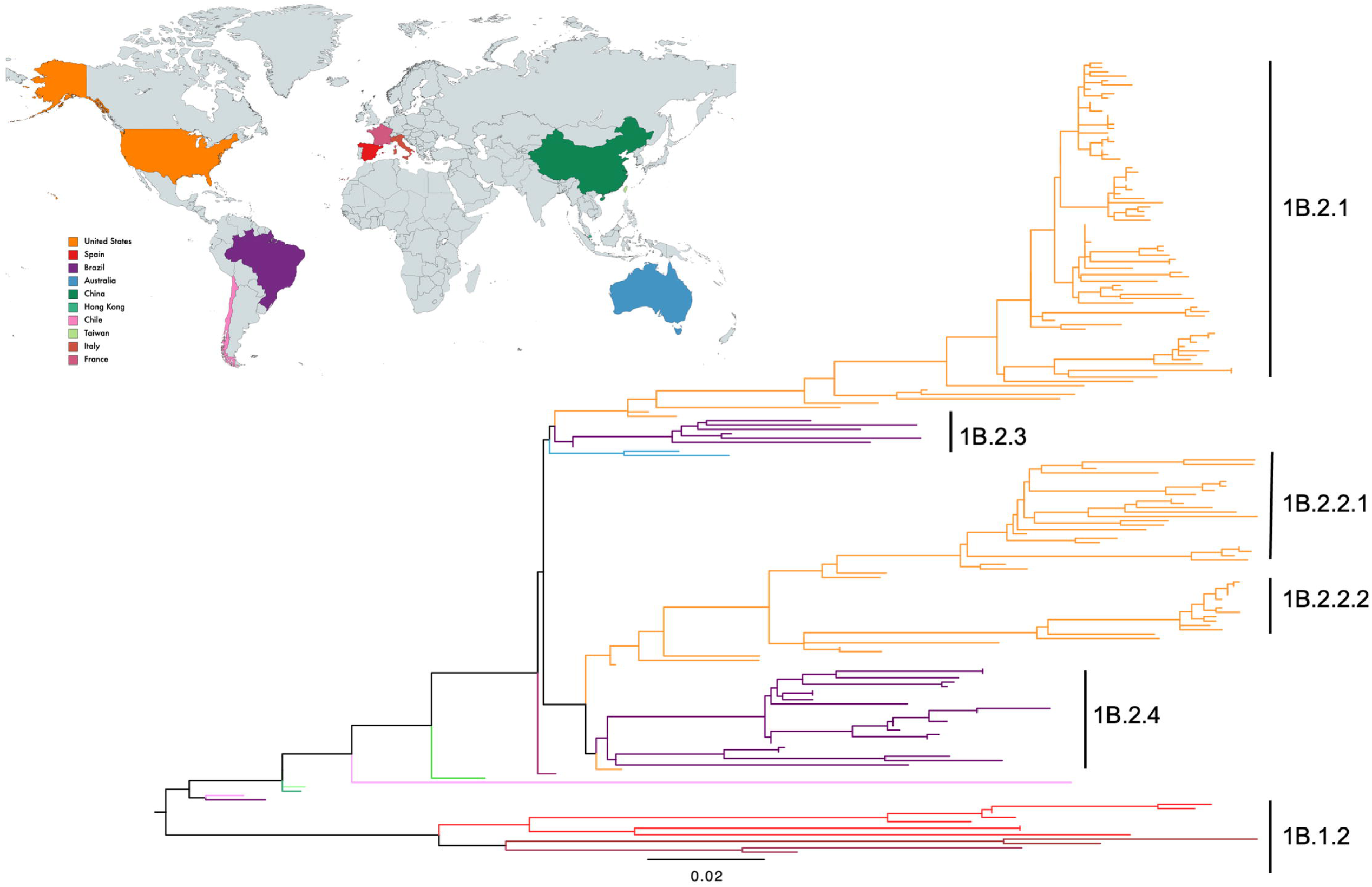
Phylogeny and geographic distribution of H1 1B human-seasonal lineage influenza A virus in swine hemagglutinin genes collected in Brazil. Mid-point rooted maximum-likelihood tree inferred for Brazilian swine HA gene sequences detected and isolated between 2010 and 2018 and global swine isolates, human seasonal strains, human vaccine, and human variant HA genes. The tree branches were colored by geographic isolation of the different strains and Brazilian HA genes were colored purple and the geographical map was generated using MapChart.

**Figure 3.**
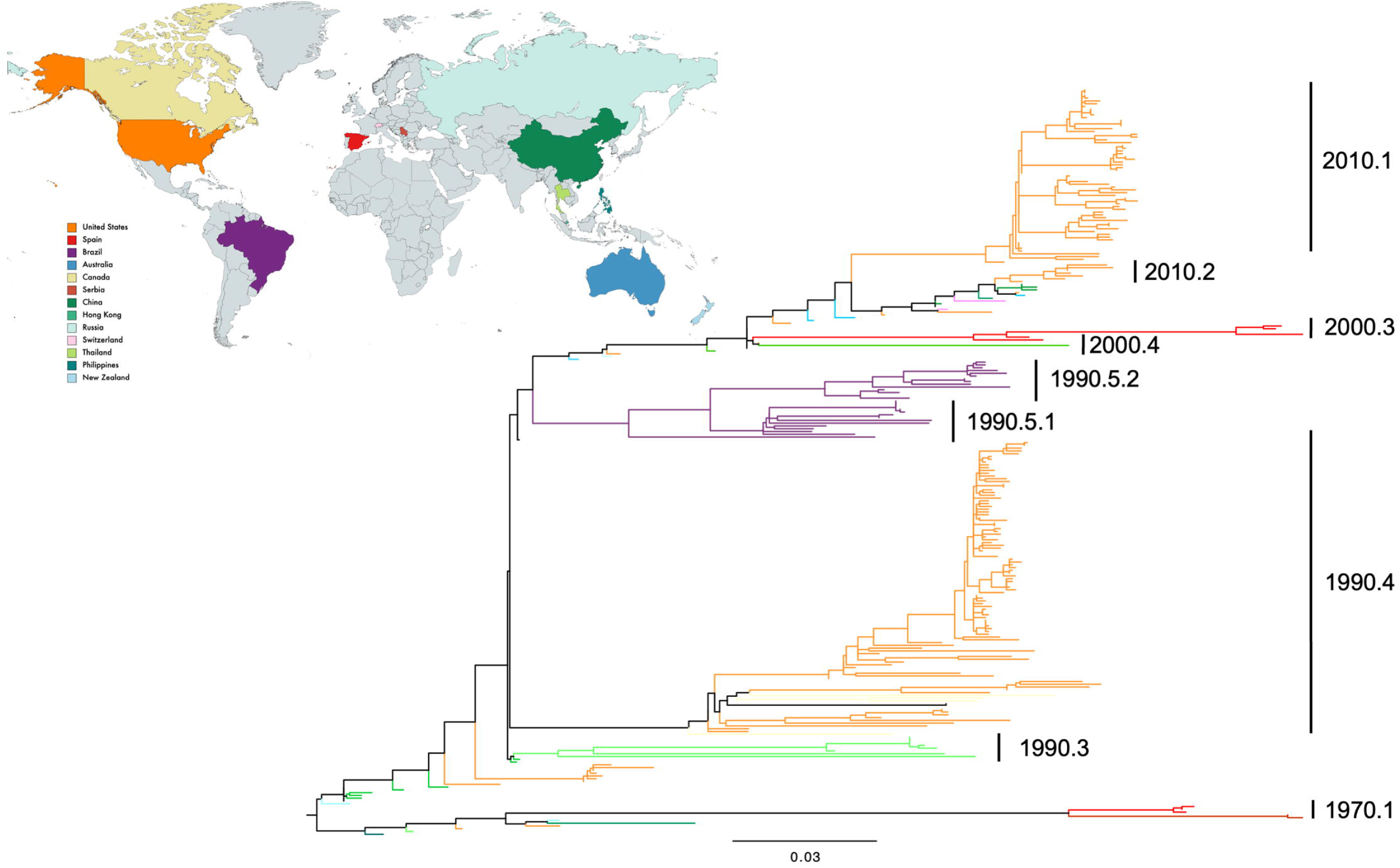
Phylogeny and geographic distribution of H3 lineage influenza A virus in swine hemagglutinin genes collected in Brazil. Mid-point rooted maximum-likelihood tree inferred for Brazilian swine HA gene sequences detected and isolated between 2010 and 2018 and global swine isolates, human seasonal strains, human vaccine, and human variant HA genes. The tree branches were colored by geographic isolation of the different strains and Brazilian HA genes were colored purple and the geographical map was generated using MapChart.

**Figure 4.**
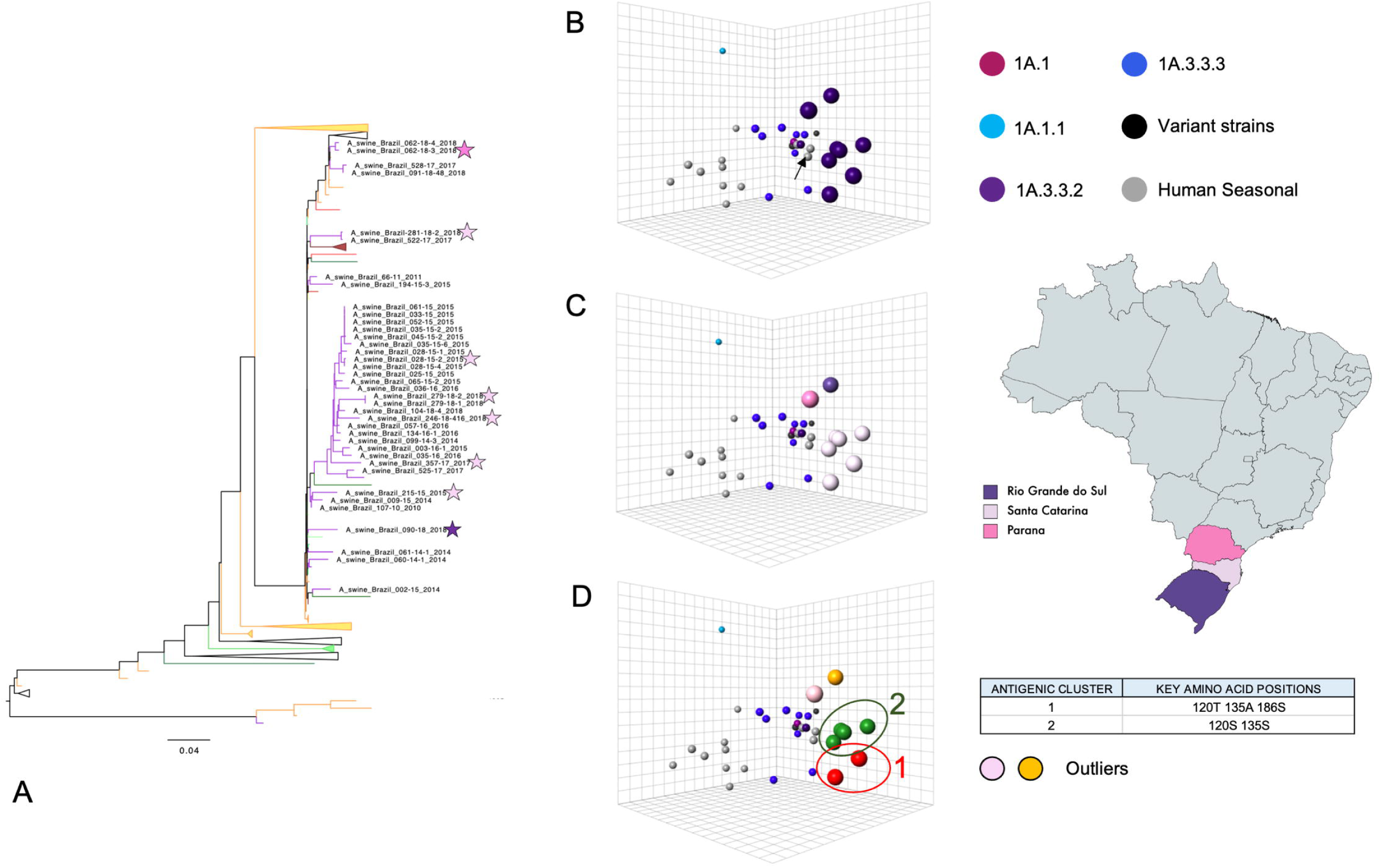
Antigenic inter-relationship between Brazilian swine H1 1A and global swine and human IAVs. (A) Mid-point rooted maximum-likelihood phylogenetic tree of Brazilian H1 1A sequences and reference global swine and human HA genes. Phylogenetic branches were coloured by virus isolation location. Clades that did not contain Brazilian isolates were collapsed for visualization. The Brazilian strains chosen for hemagglutination inhibition assays were indicated with a star colored according to isolation location. Antigenic maps were generated and the Brazilian strains were colored by genetic lineage (B), by Brazilian state (C), and by antigenic cluster (D). Brazilian strains were represented by larger dots in the antigenic map. The human seasonal vaccine strain CVV A/Michigan/45/2015 was indicated by an arrow (B). The spacing between grid lines is 1 unit of antigenic distance corresponding to a two-fold dilution of antiserum in the HI assay.

**Figure 5.**
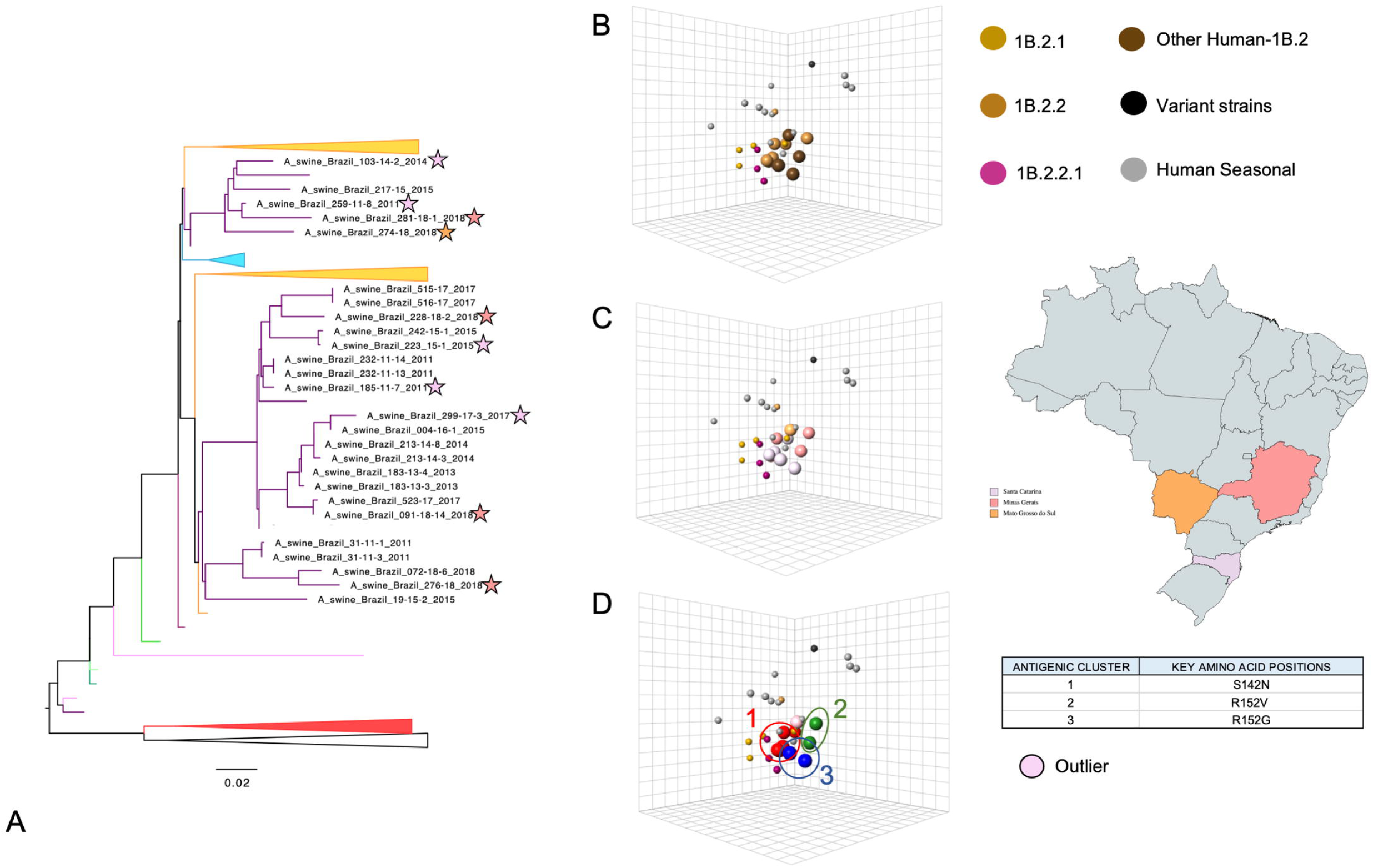
Antigenic inter-relationship between Brazilian swine H1 1B and global swine and human IAVs. (A) Mid-point rooted maximum-likelihood phylogenetic tree of Brazilian H1 1B sequences and global swine and human HA genes. Phylogenetic branches were coloured by virus isolation location. Clades that did not contain Brazilian isolates were collapsed for visualization. The Brazilian strains chosen for hemagglutination inhibition assays were indicated with a star colored according to isolation location. Antigenic maps were generated and the Brazilian strains were colored by genetic lineage (B), by Brazilian state (C), and by antigenic cluster (D). Brazilian strains were represented by the larger dots in the antigenic map. The spacing between grid lines is 1 unit of antigenic distance corresponding to a two-fold dilution of antiserum in the HI assay.

The 23 H3N2 strains circulating in Brazil were the result of a transmission event from human-to-swine in the mid-1990s (8). Within this lineage, the genetic divergence, statistical support, and evidence of onward transmission met the criteria for the naming of two clades, 1990.5.1 and 1990.5.2 (Figure 3) (following Anderson et al. (1, 21)). Rio Grande do Sul, Santa Catarina, and Mato Grosso do Sul recorded the earliest detections of 1990.5.1 strains in 2011. Since then, all H3N2 strains were isolated in the South (Santa Catarina, Parana and Rio Grande do Sul) and Midwest (Mato Grosso and Mato Grosso do Sul) of Brazil until 2016, when 1990.5.1 viruses were detected in Minas Gerais (Figure 6). Additionally, 1990.5.2 viruses were detected in Santa Catarina (2015, 2017, 2018) and Parana (2018).

**Figure 6.**
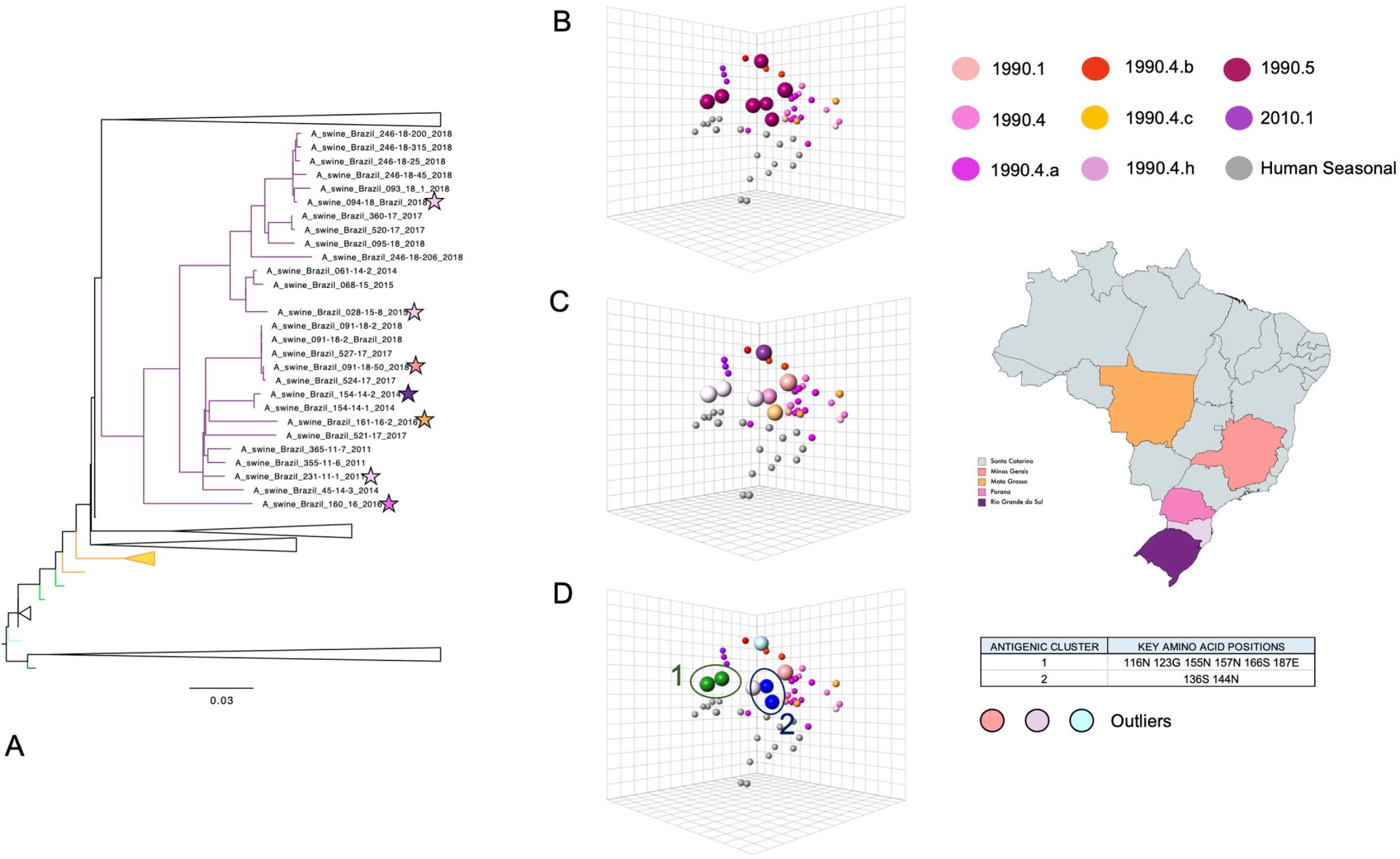
Antigenic inter-relationship between Brazilian swine H3 and global swine and human IAVs. (A) Mid-point rooted maximum-likelihood phylogenetic tree of Brazilian H3 sequences and global swine and human HA genes. Phylogenetic branches were coloured by virus isolation location. Clades that did not contain Brazilian isolates were collapsed for visualization. The Brazilian strains chosen for hemagglutination inhibition assays were indicated with a star colored according to isolation location. Antigenic maps were generated and the Brazilian strains were colored by genetic lineage (B), by Brazilian state (C), and by antigenic cluster (D). Brazilian strains were represented by the larger dots in the antigenic map. The spacing between grid lines is 1 unit of antigenic distance corresponding to a two-fold dilution of antiserum in the HI assay.

### 3.2 Antigenic characterization of swine influenza A viruses in Brazilian pigs

To quantify the antigenic evolution of 1A and 1B swine lineages in Brazil, we tested viruses isolated from clinical diagnostic submissions with swine antisera raised against reference global influenza A strains in HI assays and analysed our data using antigenic cartography. We coloured the antigenic maps by genetic lineage and clade, by geography, and by antigenic cluster. For the 1A.3.3.2 viruses, four viruses formed an antigenic cluster (green) which corresponded to a human-to-swine spillover of 1A.3.3.2 viruses that was maintained in pigs in Santa Catarina since 2015 (Figure 4). Two 1A.3.3.2 viruses detected in pigs in Santa Catarina resulted from discrete human-to-swine introductions, and were genetically distinct from each other, and also antigenically distinct from the larger Santa Catarina “green” antigenic cluster. Despite being the result of separate introductions and having 9 amino acid differences between them (four of which are in putative antigenic sites) (Table 1), A/swine/Brazil/215-15/2015 was antigenically similar to A/swine/Brazil/281-18-2/2018 (2 AU apart) and formed a second antigenic cluster (red). The other two 1A viruses from 2018 were not in a single monophyletic clade on the phylogeny, were detected in different states (Parana and Rio Grande do Sul) and were also antigenically distinct from each other and from the main ‘green’ cluster.

**Table 1.** Amino acid substitutions between Brazil H1 1A swine viruses compared to the CVV A/Michigan/45/2015 strain. Brazilian isolates have been coloured according to their antigenic clustering. Amino acid differences were calculated with Flutile python script. Antigenic units (AU) distance between human candidate vaccine (CVV) A/Michigan/45/2015 and the Brazilian 1A isolates have been added. 1 AU is equivalent to a two-fold difference in the HI assay.

| Site | REF: A/Michigan/45/2015 | A/swine/Brazil/215_15/2015 | A/swine/Brazil/281_18_2/2018 | A/swine/Brazil/028_15-2/2015 | A/swine/Brazil/357_17/2017 | A/swine/Brazil/279_18_2/2018 | A/swine/Brazil/246_18_416/2018 | A/swine/Brazil/090_18/2018 | A/swine/Brazil/062_18_3/2018 | Annotations |
| --- | --- | --- | --- | --- | --- | --- | --- | --- | --- | --- |
| AU |  | 3.5 | 3.3 | 4.9 | 4 | 3.5 | 3 | 5.1 | 3.4 |  |
| 113 | R | M |  |  |  |  |  |  |  |  |
| 120 | T |  |  | S | S | S | S |  |  |  |
| 125 | N |  | S |  |  |  | H |  |  | Sa |
| 130 | K | N |  | N | S | N | N |  |  |  |
| 135 | A |  |  | S | S | S | S |  |  |  |
| 138 | H |  |  |  |  |  |  |  | L |  |
| 141 | A |  |  |  | S |  |  | T | E |  |
| 142 | K |  | R | S |  | N | N |  |  | Ca2 |
| 146 | K |  |  |  |  |  |  | Q |  |  |
| 152 | V |  |  | I | T | I | I | I |  |  |
| 155 | G |  | E |  |  |  |  |  |  | Sa |
| 161 | L |  |  |  | F |  |  |  |  |  |
| 162 | N | S |  |  | R | S | S | I |  | Sa |
| 163 | Q | K | K | K | K | K | K | K |  | Sa |
| 166 | I |  |  |  |  | T |  |  |  | Ca1 |
| 172 | E |  | D |  |  |  |  |  |  |  |
| 173 | V | I |  |  |  |  |  |  |  |  |
| 183 | S | P |  | P | P | P | P |  |  |  |
| 185 | T | N | S | N | N | N | N | S |  |  |
| 186 | A |  |  | T | T | T | D |  |  |  |
| 187 | D | S |  | S | S | S | S |  |  |  |
| 189 | Q |  |  |  |  |  | R |  |  | Sb |
| 195 | A |  |  | E | V | E | E |  |  | Sb |
| 197 | A |  |  |  | T |  |  |  |  |  |

There is currently one commercially available vaccine against H1 1A IAVs in Brazil and it contains the A/California/04/2009 A(H1N1)pdm09 pandemic strain. In order to explore whether there was antigenic drift away from the vaccine strain through time in Brazilian pigs, we measured the antigenic distance between A/California/04/2009 and the recent 1A.3.3.2 2018 isolates. The Brazilian 1A.3.3.2 2018 isolates were located on average 3 AU away (range 2.45-4.38 AU) from swine vaccine strain A/California/04/2009 suggesting that this vaccine component is not a close vaccine-match for recently isolated Brazilian 1A strains.

In the phylogeny, the 1B swine viruses from Brazil were located within 1B.2.3 or 1B.2.4 clades within the lineage originating from pre-2009 human seasonal H1 viruses (Figure 2). These two genetic clades were distinct from other global 1B.2 clades that circulate in the USA (1B.2.1 and 1B.2.2). Despite strong phylogenetic support for these clades, the 1B genetic clade identity did not have a strict link with antigenic phenotype and similarity. The 1B.2.3 viruses were detected in pigs in Minas Gerais, Santa Catarina and Mato Grosso do Sul states (Figure 5C). The two viruses from Santa Catarina were antigenically similar (1.3 AU) and formed the blue cluster, whereas the two viruses from Minas Gerais (green) and Mato Grosso do Sul (pink) were antigenically distinct from each other and from the Santa Catarina strains by between 2.7 and 3 AU (Figure 5D). The green 1B.2.3 strain from Minas Gerais was antigenically closely related to a 1B.2.4 strain from the same state, and shared putative antigenically relevant amino acid substitution in the HA1 with the 1B.2.3 strain (153V). The five other 1B.2.4 strains comprised the red antigenic cluster and unlike the 1A viruses did not show spatial restriction, being detected in both Minas Gerais and Santa Catarina states.

The Brazilian H3 viruses evolved from a human-to-swine transmission event that occurred in the mid 1990s (8). Antigenically, the H3 strains did not cluster by phylogenetic clade identity or by year of isolation (Figure 6A and B). They also did not cluster by isolation location, as the two strains that were detected in Rio Grande do Sul were at 5.4 AU distance from each other (Figure 6C). Rather, there were two main antigenic clusters coloured blue and green and three outliers strains that were not within these two broad antigenic clusters (Figure 6D). The green cluster is composed of two 1990.5.2 lineage strains, at 1.5 AU distance from each other. The blue cluster consists of two 1990.5.1 lineage strains, at 2 AU distance from each other. The two main antigenic clusters are within 4-5 AU distance from each other.

### 3.3 Molecular determinants of antigenic diversity

To explore the molecular basis of the observed antigenic diversity in Brazilian pigs, we compared antigenically similar strains and identified amino acid differences between them, some of which were located in putative antigenic sites. We identified two different antigenic clusters in the Brazilian 1A viruses (Figure 4D). The strains A/swine/Brazil/215-15/2015 and A/swine/Brazil/281-18-2/2018 formed antigenic cluster 1 (red). Antigenic cluster 2 (green) was composed of four strains: A/swine/Brazil/028-15-2/2015, A/swine/Brazil/357-17/2017, A/swine/Brazil/279-18-2/2018 and A/swine/Brazil/246-18-416/2018. We identified amino acid substitutions T120S and A135S as potential drivers for antigenic change in these two clusters. We also compared the Brazilian 1A strains with human seasonal vaccine strains A/Michigan/45/2015 to generate a measure of zoonotic risk (Table 1). We found 6-8 amino acid substitutions (five were located in putative antigenic sites) against A/Michigan/45/2015 in antigenic cluster 1 and 11-14 substitutions (seven were located in putative antigenic sites) in antigenic cluster 2. Of note, amino acid substitution Q163K (antigenic site Sa) was present in all of the Brazilian 1A strains apart from A/swine/Brazil/062-18-3/2018.

We observed three different antigenic clusters within the Brazilian 1B viruses (Figure 5D). Antigenic cluster 1 (red) was composed of five 1B.2.4 strains and defined by amino acid 143N (Table 3). Antigenic cluster 2 (green) included the 1B.2.3 strain A/swine/Brazil/281-18-1/2018 and the 1B.2.4 strain A/swine/Brazil/276-18/2018. This cluster was defined by amino acid substitution 153V (antigenic site Sb). The 1B.2.4 strain A/swine/Brazil/276-18/2018 (green cluster) is antigenically different from all the other 1B.2.4 strains (red cluster) (Table 3). We did not identify the amino acid substitution S143N that defined the red cluster in the A/swine/Brazil/276-18/2018 strain. Lastly, antigenic cluster 3 (blue) was made up of two 1B.2.3 strains (A/swine/Brazil/259-11-8/2011 and A/swine/Brazil/103-14-2/2014) and we identified amino acid 153G (antigenic site Sb) as potentially associated with determining this antigenic group (Table 2). We also compared the HA1 genes from the Brazilian H1 1B.2.3 strains with the human seasonal vaccine A/Solomon Islands/3/2006 strain (Table 2). We identified three amino acid substitutions, T129V, P183S and K194T, in common between the Brazilian 1B.2.3 strains and that are not present in the putative ancestral strain. Brazilian strains belonging to the H1 1B.2.4 lineage were compared with a human seasonal strain A/New Caledonia/20/1999 and amino acid differences are shown in Table 3. We identified four amino acid substitutions in common to all the Brazilian 1B.2.4 isolates, T125N (antigenic site Sa), V175I, P183S and A190T (antigenic site Sb), and that were not present in the human seasonal strain.

**Table 2.** Amino acid substitutions between Brazil H1 1B.2.3 swine viruses compared to the clade ancestor strain A/Solomon Islands/3/2006. Brazilian isolates were coloured according to their antigenic clustering. Amino acid differences were calculated with flutile python script. Antigenic units (AU) distance between A/Solomon Islands/3/2006 and the Brazilian 1B.2.3 isolates have been added. 1 AU is equivalent to a two-fold difference in the HI assay.

| Site | REF: A/Solomon Islands/3/2006 | A/swine/Brazil/259_11-8/2011 | A/swine/Brazil/103_14_2/2014 | A/swine/Brazil/281_18_1/2018 | A/swine/Brazil/274-18/2018 | Annotations |
| --- | --- | --- | --- | --- | --- | --- |
| AU |  | 5.2 | 4.5 | 4.1 | 2.2 |  |
| 112 | E |  |  | D | K |  |
| 129 | T | V | V | V | V |  |
| 134 | A |  |  | S |  |  |
| 141 | E | K |  | K | K |  |
| 146 | K | R |  | R | R |  |
| 153 | G |  |  | V | R | Sb |
| 156 | G | D |  |  |  |  |
| 160 | N |  |  |  | S |  |
| 162 | S |  | R |  |  | Sa |
| 171 | K |  |  | R |  |  |
| 179 | V | I | I |  | I |  |
| 183 | P | S | S | S | S |  |
| 186 | G | R | R | R | E |  |
| 189 | R |  | K |  |  | Sb |
| 190 | A |  |  | T |  | Sb |
| 193 | H |  |  | Q |  | Sb |
| 194 | K | T | T | T | T |  |

**Table 3.** Amino acid substitutions between Brazil H1 1B.2.4 swine viruses compared to the clade ancestor strain A/New Caledonia/20/1999. Brazilian isolates have been coloured according to their antigenic clustering. Amino acid differences were calculated with Flutile python script. Antigenic units (AU) distance between A/New Caledonia/20/1999 and the Brazilian 1B.2.4 isolates have been added. 1 AU is equivalent to a two-fold difference in the HI assay.

| Site | REF: A/New Caledonia/20/1999 | A/swine/Brazil/185_11_7/2011 | A/swine/Brazil/223_15-1/2015 | A/swine/Brazil/299_17_3/2017 | A/swine/Brazil/228_18_2/2018 | A/swine/Brazil/091-18-14/2018 | A/swine/Brazil/276_18/2018 | Annotations |
| --- | --- | --- | --- | --- | --- | --- | --- | --- |
| AU |  | 2.8 | 3.1 | 2.3 | 2.3 | 1.9 | 1.2 |  |
| 104 | Q |  |  |  |  |  | H |  |
| 109 | S |  |  | P |  |  |  |  |
| 116 | I |  |  |  |  |  | V |  |
| 125 | T | N | N | N | N | N | N | Sa |
| 129 | V |  |  | I |  |  |  |  |
| 138 | H | Y | Y | Y |  | Y |  |  |
| 139 | N |  |  |  |  |  | D |  |
| 142 | S | G |  | R | G | P |  | Ca2 |
| 143 | S | N | N | N | N | N |  |  |
| 146 | R | K | K | K | K | K |  |  |
| 149 | L |  |  |  | I |  |  |  |
| 153 | G | R | R | K | R | R | V | Sb |
| 163 | K |  |  |  | R |  | N | Sa |
| 166 | V | T | A | T | T | T | T | Ca1 |
| 169 | K |  |  | R |  | R |  |  |
| 171 | K |  |  |  | R |  |  |  |
| 175 | V | I | I | I | I | I | I |  |
| 183 | P | S | S | S | S | S | S |  |
| 186 | G | K | E | E | E |  | R |  |
| 187 | D |  | N | N |  |  |  |  |
| 189 | R |  |  |  |  |  | K | Sb |
| 190 | A | T | T | T | T | I | T | Sb |
| 193 | H |  | Q |  | Q |  |  | Sb |
| 197 | A |  |  |  |  |  | T |  |
| 199 | V |  |  |  | I |  |  |  |

We observed two different antigenic clusters within the Brazilian H3 viruses (Figure 6D). Antigenic cluster 1 (green) was composed of A/swine/Brazil/028-15-8/2015 and A/swine/Brazil/094-18/2018. We identified six amino acids that defined this antigenic cluster: 117N, 124G, 156N, 158N, 167S and 188E (antigenic site B). Antigenic cluster 2 (blue) consisted of A/swine/Brazil/160-16/2016 and A/swine/Brazil/161-16-2/2016 strains. Amino acid substitutions 137S (antigenic site A) and 145N (antigenic site A) were present in both strains. We also compared the HA1 genes from the Brazilian H3 strains with the human seasonal vaccine strain, A/Wuhan/359/1995 (Table 4). We identified four amino acid substitutions present in all the H3 strains, T121N, A131S, A138S and K173E, none positioned in putative antigenic sites of the HA gene.

**Table 4.**
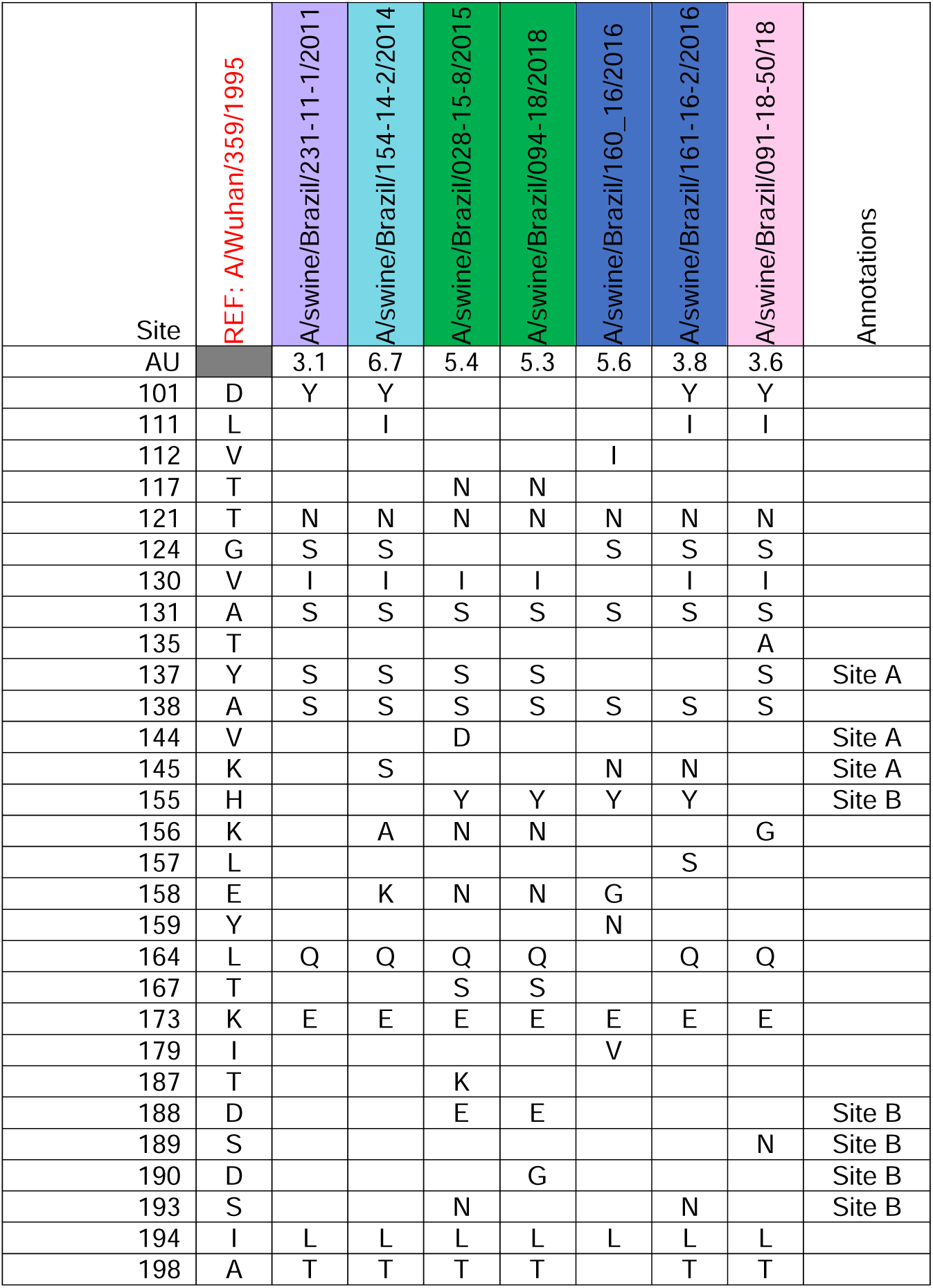
Amino acid substitutions between Brazil H3 swine viruses compared to the clade ancestor strain A/Wuhan/359/1995. Brazilian isolates were coloured according to their antigenic clustering. Amino acid differences were calculated with Flutile python script. Antigenic units (AU) distance between A/New Caledonia/20/1999 and the Brazilian 1B.2.4 isolates reported in the first row. 1 AU is equivalent to a two-fold difference in the HI assay.

### 3.4 Zoonotic risk assessment of influenza A viruses in Brazilian swine

Two human variant detections within the 1A.3.3.2 and a 1B lineage swine virus occurred in Brazil as a result of swine-to-human infection with viruses circulating in Brazilian pigs (33). Here we assessed the antigenic relationships among the Brazil H1 1A.3.3.2 lineage and sera from vaccinated swine with previous and current human seasonal H1 vaccine strains as a surrogate for the human population immunity against the swine strains due to vaccination or exposure to human seasonal viruses (Figure 7A). From the antigenic map distances, the Brazilian 1A.3.3.2 strains were antigenically greater than 5 AU from any pre-2009 human seasonal strain or from two variant human viruses (A/Minnesota/46/2015 (1A.3.3.3) and A/Ohio/09/2015 (1A.3.3.3)), to which a candidate vaccine virus (CVV) was developed. The antigenic distances to within-clade 2009-like human seasonal vaccine were smaller – averaging 3 AU away (range 3.01-3.34 AU).

**Figure 7.**
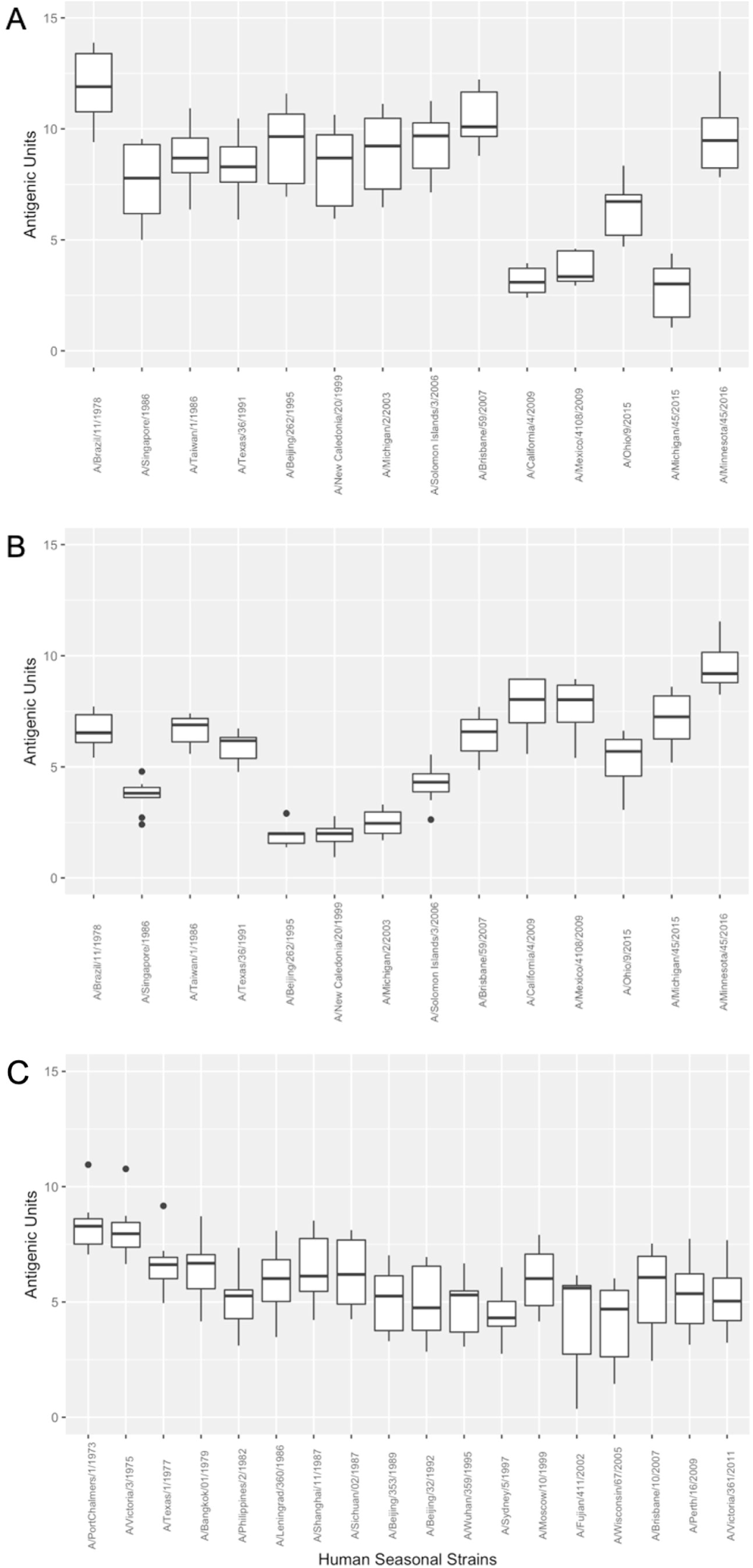
Antigenic distance between Brazilian swine H1 1A (A), 1B (B), and H3 (C) HA genes an human seasonal influenza A virus vaccines or candidate vaccine virus (CVV) strains. Antigenic unit distance between Brazil IAV in swine strains and human vaccine strains. The AU distance values were extracted from the 3D antigenic maps, and 1 unit of antigenic distance corresponds to a two-fold dilution of antiserum in the HI assay. The lines within the boxplots indicate the median value of the dataset, while the top and bottom lines correspond to the 25^th^ (Q1) and 75^th^ (Q3) quantiles. The whiskers attached to the rectangle illustrate the variability outside Q1 and Q3 quartiles.

The Brazilian 1B viruses were antigenically distant (> 5 AU) from all post-2009 human seasonal vaccine strains and to human seasonal strains that circulated between 1978 and 1991 and between 2006 and 2008 (Figure 7B). The Brazilian swine 1B viruses arose from introductions of human seasonal strains in the early 2000s and, not unexpected, we observed higher cross-reactivity between current 1B Brazilian swine viruses and human seasonal strains from 1995-2003.

We also assessed the antigenic relationships between the Brazil H3 1990.5 strains and sera from swine inoculated with previous and current human H3 vaccine strains (Figure 7C). The Brazilian H3 cluster showed a consistent antigenic distance of more than 3AU distance to all reference human seasonal H3 strains isolated between 1982-2011 (range 4.3-6.2 AU). The Brazilian H3 cluster viruses were 5.3 AU from A/Wuhan/359/1995 – a human vaccine strain in use when the Brazilian H3 lineage was established.

## 4. Discussion

In this study we characterized the genetic and antigenic diversity of H1 and H3 viruses circulating in swine in Brazil between 2010-2018. The pig samples were collected from farms located in the South, Southeast and Midwest regions of Brazil, which accounts for 92.95% of pork production in the country. The H1 and H3 tested by HI assay and mapped by antigenic cartography in this study demonstrated significant antigenic diversity. Notably, the genetic clade identity of the Brazilian swine IAV did not always strictly correlate with antigenic similarity suggesting that there may be a small number of amino acid positions that have a disproportionate impact on antigenic phenotype. Additionally, our data demonstrated that most swine strains were significantly different from the panel of human seasonal vaccine strains we tested suggesting that they may have limited efficacy in protecting human populations if these viruses were to be zoonotically transmitted.

The diversity of Brazilian H1 viruses was driven by multiple human-to-swine introductions, including the 2009 H1N1 pandemic for the Brazilian 1A lineage (1A.3.3.2) and introductions in the early 2000s for the 1B lineage (1B.2.3 and 1B.2.4) and subsequent circulation of these viruses within Brazilian pig populations (8–10). Antigenic analysis of the Brazilian 1A lineage revealed a complex spatial distribution of antigenic diversity across the lineage. One of the antigenic clusters corresponded to a single human-to-swine introduction event that has been maintained in the pigs in the state of Santa Catarina for 4 years, while the other main antigenic cluster consisted of two strains that are genetically distinct from each other but also circulate in the same region. Two other antigenically distinct 1A.3.3.2 viruses were also isolated in the states of Rio Grande do Sul and Parana. Antigenic analysis of the Brazilian 1A.3.3.2 strains revealed the formation of two main antigenic clusters with different phenotypic evolution. The ‘green’ antigenic cluster (Figure 4D) corresponded to a human-to-swine introduction that resulted in circulation and evolution in pigs while maintaining antigenic similarity; however, the second ‘red’ antigenic cluster (Figure 4D) was composed of two strains that were antigenically similar (2 AU apart) but genetically diverse (four amino acid substitutions in putative antigenic sites).

Within the 1B lineage we identified tree topologies with very long branch lengths that suggest significant unsampled diversity. There was a demonstrated increase in detections and diversity within the 1B lineage that reflects enhanced surveillance in the southeast and midwest regions of Brazil since 2017. We also observed different spatial distribution between the two different genetic clades of the Brazilian 1B lineage. The 1B.2.3 viruses were detected across Santa Catarina, Minas Gerais and Mato Grosso do Sul, forming their own distinct antigenic clusters with spatial restriction and sharing putative amino acid substitutions (153G); however, the 1B.2.4 viruses circulated in Santa Catarina and Minas Gerais and formed their own single antigenic cluster, while also defined by a single putative amino acid substitution (143N). The analysis of the Brazilian 1B viruses also demonstrated a result that was consistent across all our data: genetic clade identity was not always indicative of antigenic similarity. In this case, two strains from the two different genetic clades (1B.2.3 and 1B.2.4) were antigenically similar and even shared putative cluster-defining amino acid substitutions (153V). This was likely the result of shared evolutionary history as both of these clades of viruses were introduced into Brazilian swine in the early 2000s. Subsequent to introduction, these viruses took divergent evolutionary trajectories but this was not reflected in antigenic change, i.e., parallel evolution may have occurred as the viruses responded to new selection pressures in the new host species (34). Further surveillance may identify the maintenance of similar strains in the states of Parana and Rio Grande do Sul, perhaps with additional antigenic clusters in these pig populations.

The Brazilian H3 lineage was established in the mid 1990s as a result of a human-to-swine transmission event that has since diversified into two clades with evidence of onward transmission (1990.5.1 and 1990.5.2). The Brazilian H3 viruses were first detected in Rio Grande do Sul, Santa Catarina and Mato Grosso do Sul states in 2011. The H3 lineage viruses were also detected in the southern, southeast and midwest states of Brazil until 2018. Mato Grosso state has a single detection of H3 lineage viruses in 2016. Antigenic clustering of the Brazilian H3 viruses did not strictly follow phylogenetic relationships and although we observed two main antigenic clusters, the strains did not cluster by spatial distribution. The two main antigenic clusters were formed by strains isolated from different states in Brazil and we identified three amino acid subtitutions within putative antigenic sites (137S, 145N and 188E) that may explain this antigenic clustering.

We revealed uncharacterized antigenic diversity within IAV circulating in Brazilian swine populations and between different regions. The implementation of a single vaccine that could cover all the observed genetic and antigenic diversity would be challenging. Our data revealed five different genetic clades within three different evolutionary lineages, and these resulted in seven different antigenic clusters. Though it is potentially feasible to include seven components in a multivalent vaccine, an alternate strategy would be to conduct regular surveillance and characterization to optimize vaccine design, where vaccine strains would be selected according to the most prevalent or current antigenic cluster in circulation, rather than by spanning all diversity over a longer timeframe (35). Vaccination is currently the most cost-effective method to mitigate influenza infection in pig populations and works by promoting the generation of highly cross-protective anti-HA and anti-NA antibodies mainly using whole inactivated virus (WIV) vaccines (11, 36). However, efficacy depends on the matching of the vaccine strain to circulating IAV. In order to accurately match a vaccine strain to the circulating strains it is necessary to quantify and understand the genetic and antigenic diversity present in that region and that can be achieved through robust passive surveillance, or surveillance targeting specific populations or regions.

Surveillance and characterization of the genetic and antigenic diversity of swine IAVs in Brazil has been increasing since the H1N1 pandemic of 2009 (6, 8–10, 37). Emergence of genetically and antigenically diverse viruses in swine represents a challenge for pandemic risk preparedness. Indeed, the recent detection of swine-origin zoonotic cases in Brazil (33) highlights the importance of testing whether the current pandemic preparedness vaccines (CVVs) and human seasonal vaccines adequately capture and protect against IAV circulating in Brazilian pigs. As a baseline, human IAV vaccine strains are updated when they are 3 AU away (equivalent to 8-fold HI difference) from a circulating strain (38). Our antigenic analysis of Brazilian 1A.3.3.2 lineage reveals that all viruses were at least 3 AU away from all human seasonal vaccines. This suggests that the tested human vaccines are unlikely to protect against current 1A viruses in Brazilian pigs. 1B viruses were positioned 5 AU away from post-2009 human seasonal vaccines, and this distance is great enough that prior-exposure or vaccination with pre-2009 human seasonal vaccines will have limited protection against this lineage of swine viruses. The Brazilian H3 viruses were also positioned at least 3 AU away from all reference human seasonal H3 strains and 5 AU away from the clade ancestor strain A/Wuhan/359/1995 and more recent strains. Our results suggest that none of human vaccine strains analysed in our study are antigenically similar enough to provide significant protection against potential zoonotic infections by Brazilian H1 or H3 swine IAV.

Improved genomic surveillance of swine IAV in Brazil has allowed the characterization of representatives of circulating swine IAVs in Brazilian pigs. Our genetic and antigenic analysis of H1 and H3 viruses isolated between 2010 and 2018 revealed substantial diversity that may impact immune protection and vaccination in animals and revealed limitations in pandemic preparedness. Surveillance of swine IAVs in Brazil must continue to be a priority, especially due to the level of pork production and export that takes place in the country, which provides a greater opportunity for animal-human interaction, increasing the risk of transmission, evolution and spread of these viruses. Ongoing surveillance and genetic and antigenic characterization of circulating swine IAVs in Brazil will minimize the opportunity for vaccine failure and potential zoonotic risk in the region as well as improving control measures that can lessen the impact of this virus in animal and public health.

## Acknowledgments

We gratefully acknowledge pork producers, swine veterinarians, and laboratories for participating and publicly sharing sequences.

## Data availability statement

All Brazilian sequence data are publicly available in NCBI GenBank or GISAID with accession numbers provided in Supplementary Table XX. Data and tree files used in analyses are archived at https://github.com/flu-crew/datasets

## References

1. Anderson TK, Chang J, Arendsee ZW, Venkatesh D, Souza CK, Kimble JB, et al. Swine Influenza A Viruses and the Tangled Relationship with Humans. Cold Spring Harb Perspect Med. 2020.

2. Lewis NS, Russell CA, Langat P, Anderson TK, Berger K, Bielejec F, et al. The global antigenic diversity of swine influenza A viruses. Elife. 2016;5:e12217.

3. Chowell G, Echevarria-Zuno S, Viboud C, Simonsen L, Tamerius J, Miller MA, et al. Characterizing the epidemiology of the 2009 influenza A/H1N1 pandemic in Mexico. PLoS Med. 2011;8(5):e1000436.

4. Peiris JS, Poon LL, Guan Y. Emergence of a novel swine-origin influenza A virus (S-OIV) H1N1 virus in humans. J Clin Virol. 2009;45(3):169–73.

5. Smith GJ, Vijaykrishna D, Bahl J, Lycett SJ, Worobey M, Pybus OG, et al. Origins and evolutionary genomics of the 2009 swine-origin H1N1 influenza A epidemic. Nature. 2009;459(7250):1122–5.

6. Schaefer R, Zanella JRC, Brentano L, Vincent AL, Ritterbusch GA, Silveira S, et al. Isolation and characterization of a pandemic H1N1 influenza virus in pigs in Brazil. Pesqui Vet Brasil. 2011;31(9):761–7.

7. Caron L, Joineau MEG, Santin E, Richartz RRTB, Patrício MAdC, Soccol VT. Seroprevalence of H3N2 influenza A virus in pigs from Paraná (South Brazil): Interference of the animal management and climatic conditions. Virus Reviews & Research. 2010;15(1):3.

8. Nelson MI, Schaefer R, Gava D, Cantao ME, Ciacci-Zanella JR. Influenza A Viruses of Human Origin in Swine, Brazil. Emerg Infect Dis. 2015;21(8):1339–47.

9. Tochetto C, Junqueira DM, Anderson TK, Gava D, Haach V, Cantao ME, et al. Introductions of Human-Origin Seasonal H3N2, H1N2 and Pre-2009 H1N1 Influenza Viruses to Swine in Brazil. Viruses-Basel. 2023;15(2).

10. Junqueira DM, Tochetto C, Anderson TK, Gava D, Haach V, Cantao ME, et al. Human-to-swine introductions and onward transmission of 2009 H1N1 pandemic influenza viruses in Brazil. Frontiers in Microbiology. 2023;14.

11. Vincent AL, Perez DR, Rajao D, Anderson TK, Abente EJ, Walia RR, et al. Influenza A virus vaccines for swine. Vet Microbiol. 2017;206:35–44.

12. Vincent AL, Anderson TK, Lager KM. A Brief Introduction to Influenza A Virus in Swine. Methods Mol Biol. 2020;2123:249–71.

13. Zhang J, Harmon KM. RNA Extraction from Swine Samples and Detection of Influenza A Virus in Swine by Real-Time RT-PCR. Methods Mol Biol. 2020;2123:295–310.

14. Zhang J, Gauger PC. Isolation of Swine Influenza A Virus in Cell Cultures and Embryonated Chicken Eggs. Methods Mol Biol. 2020;2123:281–94.

15. Zhang Y, Aevermann BD, Anderson TK, Burke DF, Dauphin G, Gu Z, et al. Influenza Research Database: An integrated bioinformatics resource for influenza virus research. Nucleic Acids Res. 2017;45(D1):D466–D74.

16. Katoh K, Standley DM. MAFFT multiple sequence alignment software version 7: improvements in performance and usability. Molecular biology and evolution. 2013;30(4):772–80.

17. Nguyen LT, Schmidt HA, von Haeseler A, Minh BQ. IQ-TREE: a fast and effective stochastic algorithm for estimating maximum-likelihood phylogenies. Mol Biol Evol. 2015;32(1):268–74.

18. Kalyaanamoorthy S, Minh BQ, Wong TKF, von Haeseler A, Jermiin LS. ModelFinder: fast model selection for accurate phylogenetic estimates. Nat Methods. 2017;14(6):587–9.

19. Guindon S, Dufayard JF, Lefort V, Anisimova M, Hordijk W, Gascuel O. New algorithms and methods to estimate maximum-likelihood phylogenies: assessing the performance of PhyML 3.0. Syst Biol. 2010;59(3):307–21.

20. Hoang DT, Chernomor O, von Haeseler A, Minh BQ, Vinh LS. UFBoot2: Improving the Ultrafast Bootstrap Approximation. Mol Biol Evol. 2018;35(2):518–22.

21. Anderson TK, Macken CA, Lewis NS, Scheuermann RH, Van Reeth K, Brown IH, et al. A Phylogeny-Based Global Nomenclature System and Automated Annotation Tool for H1 Hemagglutinin Genes from Swine Influenza A Viruses. mSphere. 2016;1(6):e00275–16.

22. Chang J, Anderson TK, Zeller MA, Gauger PC, Vincent AL. octoFLU: Automated Classification for the Evolutionary Origin of Influenza A Virus Gene Sequences Detected in US Swine. Microbiology resource announcements. 2019;8(32):e00673–19.

23. Burke DF, Smith DJ. A recommended numbering scheme for influenza A HA subtypes. PLoS One. 2014;9(11):e112302.

24. Wiley DC, Wilson IA, Skehel JJ. Structural identification of the antibody-binding sites of Hong Kong influenza haemagglutinin and their involvement in antigenic variation. Nature. 1981;289(5796):373–8.

25. Caton AJ, Brownlee GG, Yewdell JW, Gerhard W. The antigenic structure of the influenza virus A/PR/8/34 hemagglutinin (H1 subtype). Cell. 1982;31(2 Pt 1):417–27.

26. Kitikoon P, Gauger PC, Vincent AL. Hemagglutinin inhibition assay with swine sera. Methods Mol Biol. 2014;1161:295–301.

27. Lewis NS, Anderson TK, Kitikoon P, Skepner E, Burke DF, Vincent AL. Substitutions near the hemagglutinin receptor-binding site determine the antigenic evolution of influenza A H3N2 viruses in U.S. swine. J Virol. 2014;88(9):4752–63.

28. Rajao DS, Anderson TK, Kitikoon P, Stratton J, Lewis NS, Vincent AL. Antigenic and genetic evolution of contemporary swine H1 influenza viruses in the United States. Virology. 2018;518:45–54.

29. Bolton MJ, Abente EJ, Venkatesh D, Stratton JA, Zeller M, Anderson TK, et al. Antigenic evolution of H3N2 influenza A viruses in swine in the United States from 2012 to 2016. Influenza Other Respir Viruses. 2019;13(1):83–90.

30. Smith DJ, Lapedes AS, de Jong JC, Bestebroer TM, Rimmelzwaan GF, Osterhaus AD, et al. Mapping the antigenic and genetic evolution of influenza virus. Science. 2004;305(5682):371–6.

31. Team RC. R: A language and environment for statistical computing. 2015.

32. Abente EJ, Santos J, Lewis NS, Gauger PC, Stratton J, Skepner E, et al. The molecular determinants of antibody recognition and antigenic drift in the H3 hemagglutinin of swine influenza A virus. Journal of virology. 2016;90(18):8266–80.

33. Resende PC, Born PS, Matos AR, Motta FC, Caetano BC, Debur MD, et al. Whole-Genome Characterization of a Novel Human Influenza A(H1N2) Virus Variant, Brazil. Emerg Infect Dis. 2017;23(1):152–4.

34. Xiang D, Shen X, Pu Z, Irwin DM, Liao M, Shen Y. Convergent Evolution of Human-Isolated H7N9 Avian Influenza A Viruses. J Infect Dis. 2018;217(11):1699–707.

35. Walia RR, Anderson TK, Vincent AL. Regional patterns of genetic diversity in swine influenza A viruses in the United States from 2010 to 2016. Influenza Other Respir Viruses. 2019;13(3):262–73.

36. Ellebedy AH, Webby RJ. Influenza vaccines. Vaccine. 2009;27 Suppl 4(Suppl 4):D65–8.

37. Rajao DS, Costa AT, Brasil BS, Del Puerto HL, Oliveira FG, Alves F, et al. Genetic characterization of influenza virus circulating in Brazilian pigs during 2009 and 2010 reveals a high prevalence of the pandemic H1N1 subtype. Influenza Other Respir Viruses. 2013;7(5):783–90.

38. Ampofo WK, Azziz-Baumgartner E, Bashir U, Cox NJ, Fasce R, Giovanni M, et al. Strengthening the influenza vaccine virus selection and development process: Report of the 3rd WHO Informal Consultation for Improving Influenza Vaccine Virus Selection held at WHO headquarters, Geneva, Switzerland, 1-3 April 2014. Vaccine. 2015;33(36):4368–82.

