## Supplemental Table 1 for "Antigenic and genetic diversity of H1 and H3 influenza A viruses in swine in Brazil"

**Supplementary Material**

Table S1: Influenza A strains detected and isolated from pigs in Brazil between 2010-2018. Brazil isolates chosen for the HI assay are indicated by a triangle.

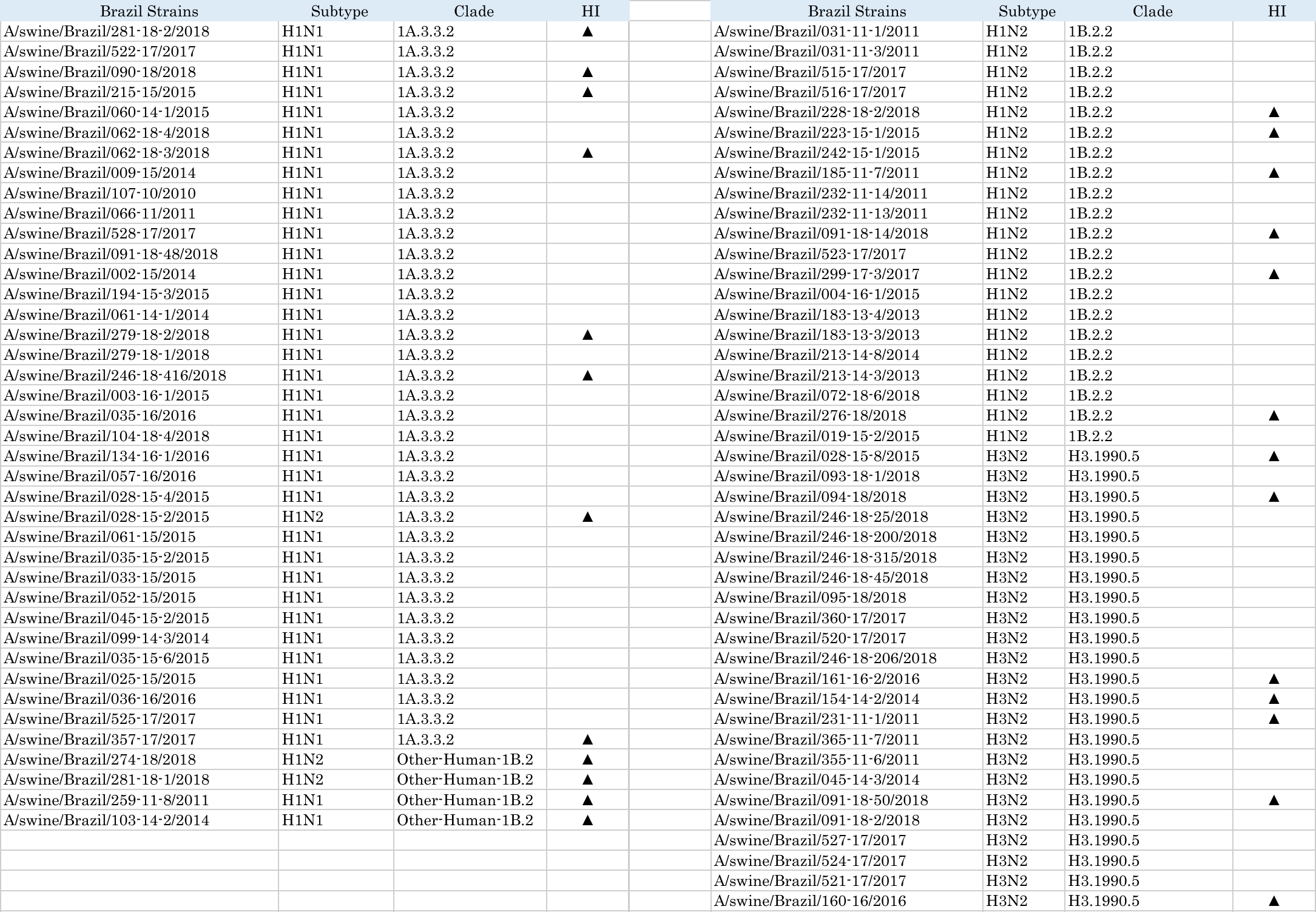
